# Chondrolectin regulates the sublaminar localization and regenerative function of muscle satellite cells in mice

**DOI:** 10.1101/2025.09.09.675211

**Authors:** Lijie Gu, Kun Ho Kim, Xiyue Chen, Stephanie N Oprescu, Yufen Li, Junxiao Ren, Shihuan Kuang, Feng Yue

## Abstract

Skeletal muscle satellite cells (SCs) reside between the myofiber sarcolemma and basal lamina, where extracellular matrix (ECM) interactions are essential for their maintenance and regenerative function. Here, we identify chondrolectin (CHODL), a type I transmembrane protein with a C-type lectin domain, as a critical regulator of SC biology. Single-cell RNA-seq analysis reveals that *Chodl* is highly enriched in quiescent SCs but downregulated in proliferating myoblasts. Using conditional knockout models, we show that deletion of *Chodl* in embryonic myoblasts (*Chodl^MKO^*) or adult SCs (*Chodl^PKO^*) does not affect muscle development but markedly impairs regeneration in both young and aged mice. *Chodl*-deficient SCs exhibit reduced self-renewal, diminish proliferation, and impair differentiation, leading to defective myofiber repair. In silico network perturbation further predicts disruption of ECM-ligand interactions and Notch signaling, consistent with our observation that a significant fraction of SCs in *Chodl^PKO^*mice localize outside the basal lamina and undergo precocious activation. Together, these findings establish CHODL as a key determinant of SC niche localization and regenerative function, uncovering a previously unrecognized mechanism linking ECM interactions to muscle stem cell maintenance and repair.

## Introduction

Tissue-resident adult stem cells possess a remarkable ability to continuously regenerate local tissues. Satellite cells (SCs), the resident stem cells of adult skeletal muscle, play a critical role in muscle regeneration. Residing beneath the basal lamina in close association with myofibers, SCs maintain a quiescent state under homeostatic conditions (Kuang et al., 2008; Yin et al., 2013). Upon muscle injury or other stimuli, SCs exit quiescence, re-enter the cell cycle, and undergo metabolic and transcriptional remodeling that enables their migration and proliferation (de Morree and Rando, 2023; Relaix et al., 2021; Sartorelli and Ciuffoli, 2025; Yue et al., 2022). These expanded SCs either fuse to repair damaged fibers or self-renew to replenish the stem cell pool (Relaix et al., 2021). The transition between these fates quiescence, activation, differentiation, and self-renewal, is tightly regulated by both various intrinsic programs and extrinsic signals from the surrounding microenvironment (Brunet et al., 2023; Fuchs and Blau, 2020; Hicks and Pyle, 2023). Niche regulation of muscle stem cell function is influenced by the composition and architecture of the extracellular matrix (ECM) (Fuchs and Blau, 2020). Following muscle injury, SCs remain encased within the residual basal lamina, referred to as a “ghost fiber”, which serves as a structural scaffold for regeneration (Vracko and Benditt, 1972; Webster et al., 2016). However, disruption of this basal lamina can lead to disorganized myofiber regeneration, misdirected SC migration, and abnormal myotube formation (Rayagiri et al., 2018; Webster et al., 2016). Recent studies reported that ECM components, including laminin, collagen, integrin, and fibronectin, etc., are key in regulating SC fate in a cell-autonomous manner (Baghdadi et al., 2018; Chang et al., 2018; Ishii et al., 2018; Lukjanenko et al., 2016; Penton et al., 2016; Rayagiri et al., 2018; Urciuolo et al., 2013). Laminin, a key component of the basal lamina, directly interacts with integrins and other SC surface receptors to regulate adhesion, polarity, and fate decisions (Penton et al., 2016; Rayagiri et al., 2018). Similarly, collagen not only provides structural support but also modulates SC behavior by influencing stiffness and mechanical signaling (Baghdadi et al., 2018; Urciuolo et al., 2013). Additionally, ECM remodeling enzymes actively reshape the basal lamina during repair, modifying the physical and biochemical environment of SCs (Goetsch et al., 2003; Thomas et al., 2015). These ECM components actively instruct SC function, highlighting the niche as a dynamic regulator of muscle regeneration, though the precise molecular mediators and underlying mechanisms remain incompletely understood.

Chondrolectin (CHODL) is a type I transmembrane protein with poorly understood cellular function (Weng et al., 2003). CHODL harbors a C-type lectin domain, which is responsible for recognizing and binding to carbohydrate, proteins with C-type lectin domain have diverse functions including cell-cell adhesion, ligand binding, and immune response to pathogens (Weis et al., 1998; Zelensky and Gready, 2005). Recent studies have shown that the extracellular domain of CHODL interacts with collagen components in the ECM and plays important roles in the regulation of motor neuron functions, promoting cell survival and neurite outgrowth (Enjin et al., 2010; Oprisoreanu et al., 2019). Using *In situ* hybridization on E15 mouse embryo, it was revealed that CHODL was highly expressed in muscle cells of various organs (Claessens et al., 2007; Weng et al., 2003). Although CHODL was shown to ameliorate motor neuron outgrowth defects in a zebrafish model of spinal muscular atrophy (Sleigh et al., 2014), its roles in SCs has not been undefined.

In this study, we analyzed published single-cell RNA-seq (scRNA-seq) datasets and found that CHODL is highly expressed in quiescent SCs compared with activated and differentiated SCs. We hypothesized that CHODL may contribute to muscle development or regeneration. To test this, we generated a conditional knockout (KO) mouse model using *MyoD^Cre^* mice to specifically delete the *Chodl* gene in embryonic myoblasts (*Chodl^MKO^* mice). While *Chodl^MKO^* mice showed normal muscle growth and myofiber formation, *Chodl* KO SCs displayed altered behaviors during muscle development. *Chodl^MKO^* muscles exhibited compromised regeneration capacity in young and aged mice. To specifically study the effect of CHODL in SCs, we crossed *Pax7^CreER^*mice with *Chodl^flox/flox^* mice and generated tamoxifen inducible SC-specific *Chodl* KO mice (*Chodl^PKO^* mice). *Chodl^PKO^* mice exhibited diminished number of SCs, increased SC detachment from myofibers, and impaired muscle regeneration. Collectively, our study demonstrates a critical role of CHODL in SC biology in regulating muscle homeostasis and regeneration.

## Results

### CHODL is abundantly expressed in quiescent SCs

To gain insight into the expression pattern of *Chodl* across various tissues, we conducted a comprehensive analysis using scRNA-seq data publicly available from *Tabula Muris* (Tabula Muris et al., 2018). Our analysis revealed that *Chodl* mRNA was mainly enriched in SCs, bladder cells and a non-annotated cell population (**Fig. 1A, B**). Moreover, we analyzed our previously published scRNA-seq data (Oprescu et al., 2020b) to pinpoint the expression of *Chodl* in subsets of myogenic cells isolated from non-injured and regenerating muscles at 5 and 10 days post injury (DPI). UMAP reveals a similar expression pattern between *Chold* and *Pax7*, moderate levels of *Chodl* in proliferating (Mik67^+^) cells, but mutually exclusive expression patterns between *Chodl* and *Myog* (**Fig. 1C**). Consistently, violin plots show that *Chodl* transcripts were predominantly detected in quiescent (QSC) and self-renewal (SSC), moderately detected in activated (ASC) and proliferating (PSC), but not in committed (CSC) and differentiating (DSC) SCs (**Fig. 1D**).

**Figure 1.**
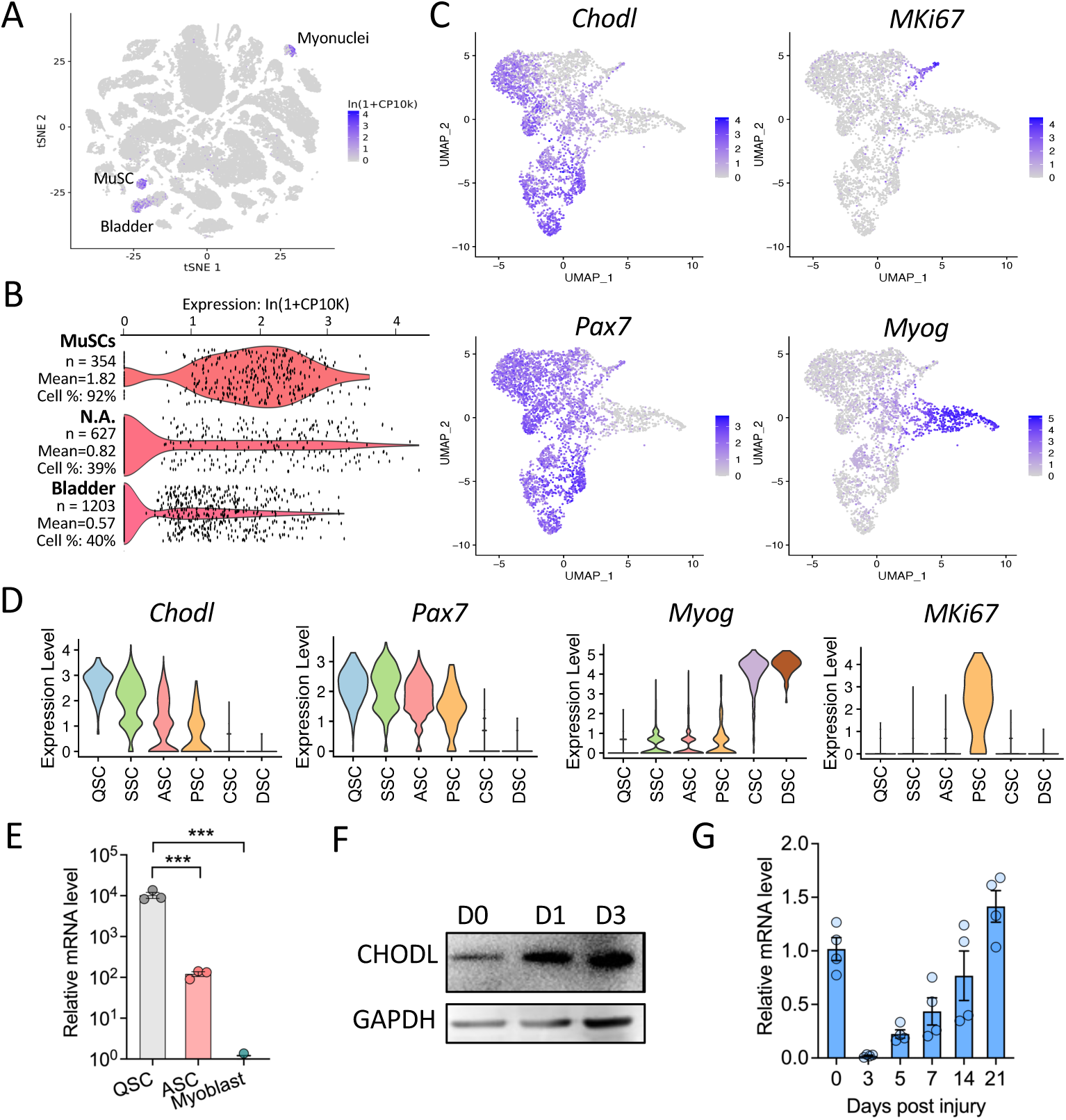
*Chodl* mRNA is highly expressed in quiescent SCs. **(A)** *Chodl* gene expression in various cell types in mouse. Data was explored through the *Tabula Muris* data portal. **(B)** Relative expression of *Chodl* gene in SCs, bladder cells and a non-annotated cell population. **(C)** UMAP-embedding of scRNA-seq data on SCs isolated from non-injured muscles and injured muscles at 5 dpi in mouse. **(D)** Violin plots showing subcluster-specific gene expression (*Pax7*, *Myog*, *Mki67*) and enrichment of *Chodl* in the QSCs and ASCs. Colored by cluster identity. Abbreviations: QSC, quiescent SCs; SSC, self-renewal SCs; ASC, activated SCs; PSC, proliferating SCs; CSC, committed SCs; DSC, differentiating SCs. **(E)** qRT-PCR analysis of *Chodl* expression level in mouse QSC, ASC and myoblast. n=3 mice. **(F)** Immunoblot analysis showing CHODL expression in mouse myoblasts during myogenic differentiation. **(G)** qRT-PCR analysis of *Chodl* in TA muscles post CTX injury in mouse. n=4 mice. Data are represented as mean ± SEM; Unpaired Student’s *t*-test; * *P*< 0.05, ** *P*< 0.01, *** *P*< 0.001.

To validate these findings, we performed qPCR analysis of *Chodl* mRNA levels among distinct SCs populations, including QSCs (isolated from uninjured muscles), ASCs (isolated from muscles 3.5 days post injury), and cultured primary myoblasts. *Chodl* was highly expressed in QSCs and moderately expressed in ASCs but nearly undetectable in cultured myoblasts (**Fig. 1E**). To further profile Chodl expression during later stages of myogenesis, we differentiated primary myoblasts for 3 days. Immunoblotting analysis indicates that CHODL protein levels increased during myoblast differentiation (**Fig. 1F**). In addition, we examined *Chodl* mRNA in skeletal muscles at various time points after muscle injury (**Fig. 1G**). We found that *Chodl* mRNA was diminished at 3 dpi, and the levels gradually returned to the uninjured level after 14 dpi. These data collectively demonstrate that *Chodl* is highly expressed in SCs and differentiated myofibers in the skeletal muscle.

### Myogenic lineage-specific ablation of *Chodl* does not affect muscle development

To directly examine the physiological role of CHODL in SCs, we generated an embryonic myogenic progenitor-specific *Chodl* KO mice (*Chodl^MKO^*) by crossing *Chodl^flox/flox^* mice with *MyoD^Cre^* mice expressing Cre recombinase driven by the endogenous *Myod1* gene promoter (**Fig. 2A**). A previous study has demonstrated that *MyoD^Cre^* selectively targets embryonic myogenic progenitors that subsequently differentiate into postnatal SC and myofibers (Kanisicak et al., 2009). In this mouse model, the deletion of exon 2-4 in the *Chodl* gene in myogenic lineage cells induces a frameshift and generates a premature stop codon, leading to the production of truncated proteins, in which the CLECT functional domain was missing (**Fig. 2A**). We confirmed the effective ablation of CHODL at both protein and mRNA levels in skeletal muscles of *Chodl^MKO^*mice (**Fig. 2B, C**).

**Figure 2.**
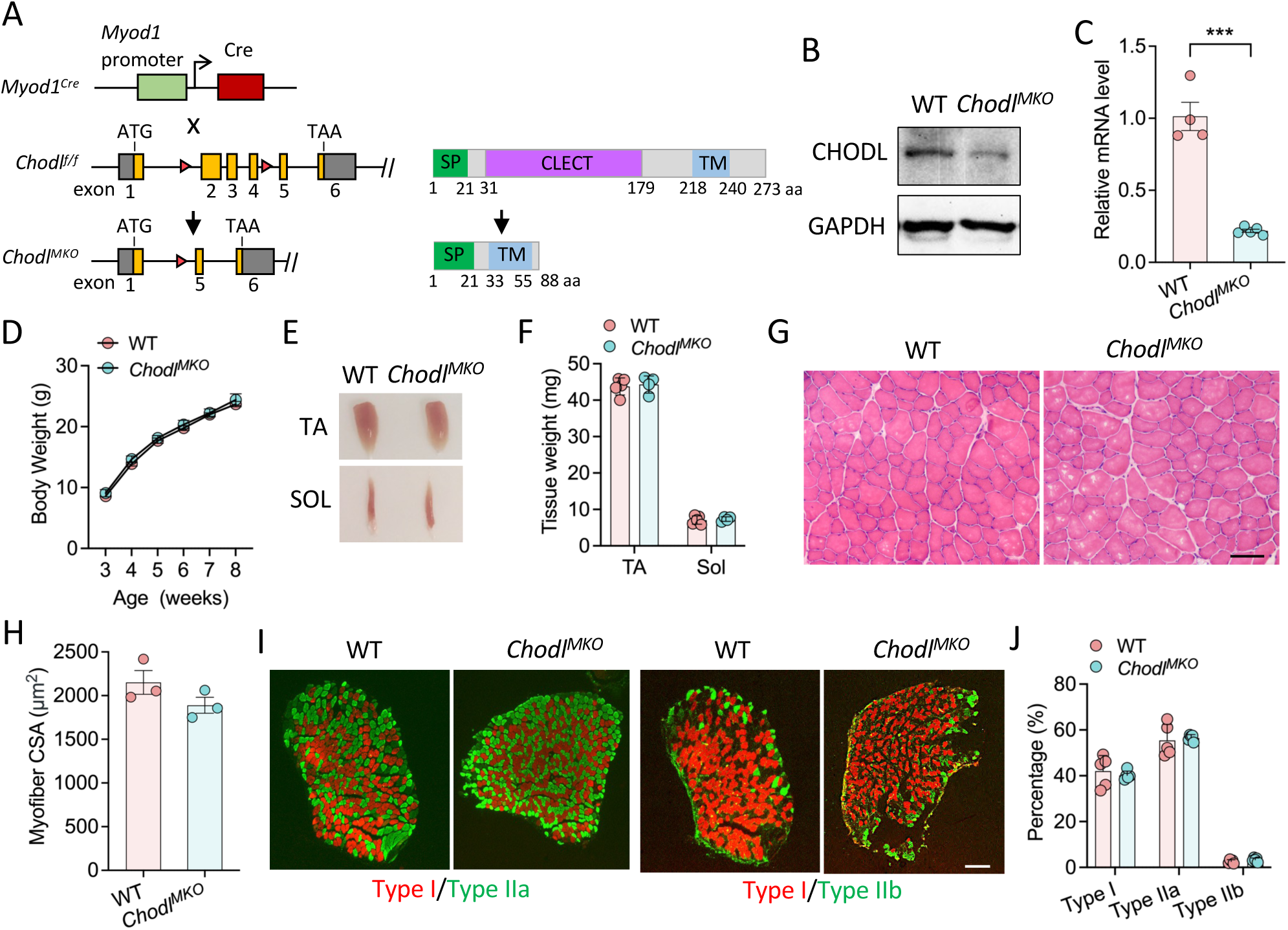
*Chodl* is dispensable for postnatal muscle development. **(A)** Gene targeting strategy for generating *Chodl^MKO^* mice. **(B, C)** Western blot (**B**) and qPCR (**C**) analysis of TA muscle samples from young adult WT or *Chodl^MKO^* mice. n=5 mice each group. **(D)** Growth curve of WT and *Chodl^MKO^* mice. n=5 mice each group. **(E)** Representative images of TA muscles and soleus muscles in young adult WT and *Chodl^MKO^* mice. **(F)** Skeletal muscle tissue weight. TA, Tibialis anterior muscle. SOL, Soleus muscle. n=5 mice each group. **(G)** H&E staining of young adult WT and *Chodl^MKO^* mice TA muscle cross-sections. Scale bar: 50 μm. **(H)** Average myofiber CSA of TA muscle section. n=3 mice each group. **(I)** Representative fiber-typing images of soleus muscles. Scale bar: 200 μm. **(J)** Abundancy of Type I, Type IIa and Type IIb myofibers in soleus muscles of WT and *Chodl^MKO^* mice. n=5 mice each group. Data are represented as mean ± SEM; Unpaired Student’s *t*-test; *** *P*< 0.001.

The *Chodl^MKO^* mice were born in expected Mendelian ratio and had the same growth rate as the WT littermates (**Fig. 2D**). The morphology and weight of skeletal muscles were indistinguishable between adult WT and *Chodl^MKO^* mice (**Fig. 2E, F**). Histologically, WT and *Chodl^MKO^* TA muscles exhibit similar myofiber size with no difference in the cross-sectional area (CSA) (**Fig. 2G, H**). As CHODL is also abundantly expressed in myofibers, we evaluated myofiber composition in the soleus muscles utilizing immunofluorescent staining of myosin heavy chain isoforms. We found that myofiber size and fiber type distribution in soleus muscles was comparable between the WT and *Chodl^MKO^*mice (**Fig. 2I, J**). Collectively, these observations suggest that CHODL in myogenic progenitors is dispensable for maintaining muscle growth and fiber composition under normal physiology.

### Loss of *Chodl* reduces muscle SC pool and impairs muscle regeneration in young adult mice

Given the high expression of *Chodl* in SCs, we sought to examine whether loss of *Chodl* alters the SC compartment in adult mice. Immunofluorescence staining of PAX7 and α-laminin revealed a significantly less PAX7^+^ SCs observed in TA muscles of *Chodl^MKO^* mice compared to WT mice at 2-month-old (**Fig. 3A**). Specifically, the number of PAX7^+^ SCs per TA area was reduced by 52% (2.3 versus 4.8 SCs per TA area) (**Fig. 3B**). These findings suggest that CHODL plays an important role in maintaining the SC pool.

**Figure 3.**
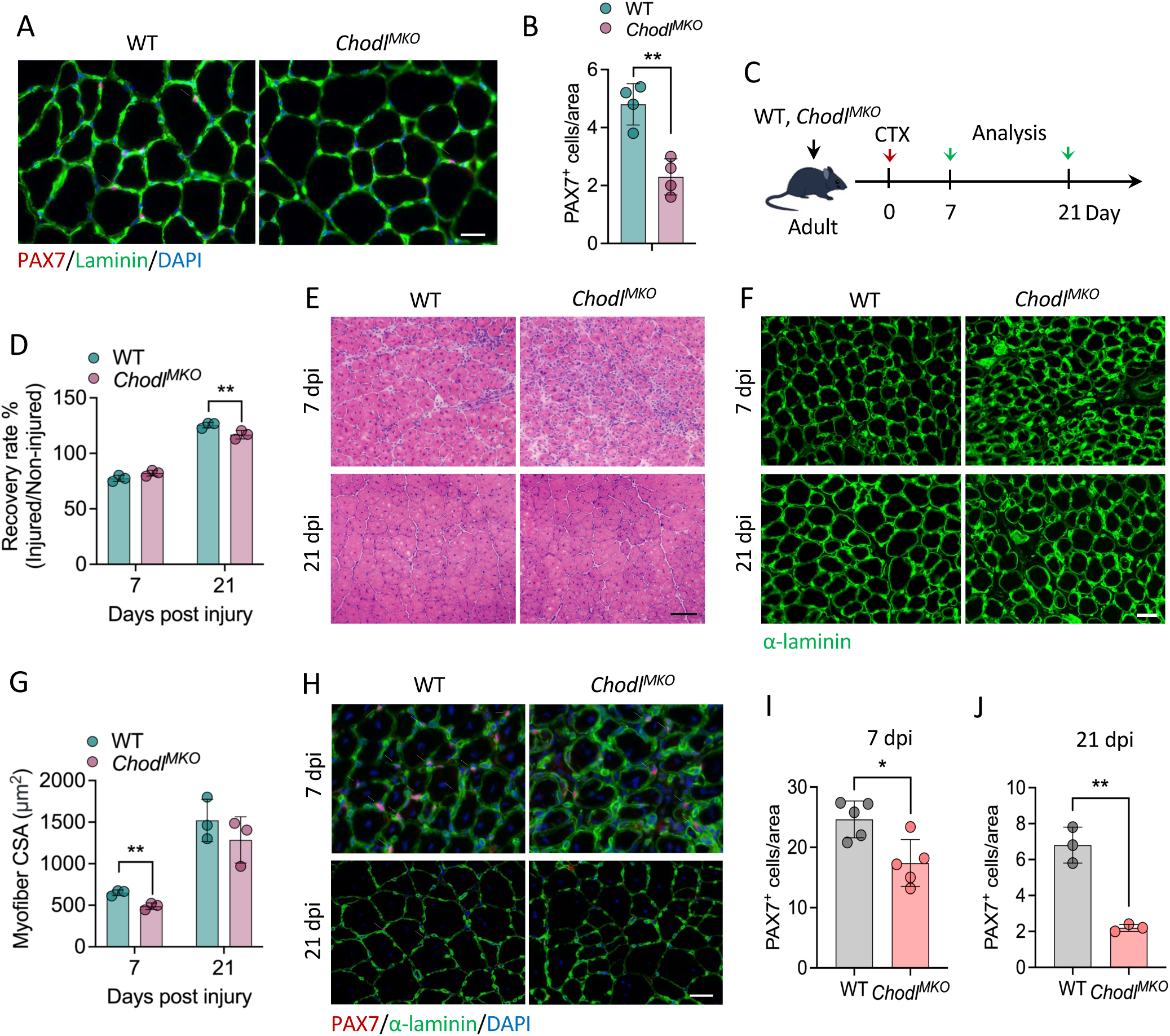
Loss of *Chodl* reduces SC number and impairs muscle regeneration. (**A**) Immunofluorescence of PAX7, α-laminin on TA muscle cross-sections from young adult WT and *Chodl^MKO^*. Scale bar: 50 μm. (**B**) Quantification of PAX7^+^ cells per area as shown in (**A**), n=4 mice each group. (**C**) Schematics showing experimental design involving cardiotoxin (CTX)-induced muscle regeneration in young adult mice (8 to 12-week-old). (**D**) TA muscle recovery rate calculated by the ratio of injured to non-injured TA muscle weights at 7 and 21 days post injury (dpi). n=3 mice each group. (**E**) H&E and immunofluorescence staining of WT and *Chodl^MKO^* mice TA muscle cross-sections at 7 and 21 dpi. Scale bar: 50 μm. n=3 each group. (**F**) Immunofluorescence of α-laminin on WT and *Chodl^MKO^* TA muscle cross-sections at 7 and 21 dpi. Scale bar: 50 μm. (**G**) Average myofiber CSA of TA muscle section. n=3 mice each group. (**H**) Immunofluorescence of PAX7, α-laminin on WT and *Chodl^MKO^*TA muscle cross-sections at 7 and 21 dpi. Scale bar: 50 μm. **(I, J)** Quantification of PAX7^+^ cell number per area at 7 (**I**) and 21 dpi (**J**). as shown in (**H**). For 7 dpi, n=5 mice each group; For 21 dpi, n=3 mice each group. Data are represented as mean ± SEM; Unpaired Student’s *t*-test; \**P*< 0.05, \*\**P* < 0.01.

To investigate the physiological function of CHODL in SCs during muscle repair, we assessed regeneration following cardiotoxin (CTX)-induced injury of tibialis anterior (TA) muscles in young adult *Chodl^MKO^* mice (**Fig. 3C**). Compared with WT controls, *Chodl^MKO^*mice exhibited a significantly reduced muscle recovery rate, as determined by the ratio of injured to uninjured TA muscle mass, at 21 days post-injury (dpi), whereas recovery was comparable to WT controls at 7 dpi (**Fig. 3D**). Histological H&E staining showed that the *Chodl^MKO^* muscles contained smaller centrally nucleated myofibers at both 7 and 21 dpi, as well as greater infiltration of interstitial mononuclear cells at 7 dpi, compared with the well-regenerated WT muscles (**Fig. 3E**). Consistently, immunofluorescence staining of α-laminin showed that *Chodl^MKO^*myofibers were significantly smaller than WT controls at 7 dpi, with a trend toward reduction at 21 dpi (**Fig. 3F, G**). These results suggest that deletion of *Chodl* in embryonic myoblasts results in the defect of muscle regeneration. We further performed immunofluorescence staining of PAX7 and found a significant reduction in the number of PAX7^+^ SCs in the *Chodl^MKO^* muscles compared to WT at 7 dpi (**Fig. 3H, I**). Remarkably, the number of PAX7^+^ cells was reduced by 70% in *Chodl^MKO^* mice at 21 dpi, when the muscle regeneration was completed and self-renewed SCs were returned to quiescent state in WT mice (**Fig. 3H, J**). Together, these findings indicate an impairment in the self-renewal and regenerative capacity of *Chodl*-deficient SCs.

### *Chodl* KO compromises SC maintenance and muscle regeneration in aged mice

Given the observation of reduced self-renewal potential of *Chodl*-deficient SCs, we next examined the number of SCs in aged mice. At 22 months of age, a significant decrease in SC abundance was observed in TA muscles of *Chodl^MKO^* mice compared to that of WT mice (**Fig. 4A, B**). Similarly, freshly isolated single myofiber from *Chodl^MKO^* mice contained fewer SCs than those from WT mice (**Fig. 4C, D**), suggesting that *Chodl*-deficient SCs fail to maintain SC pool during aging.

**Figure 4.**
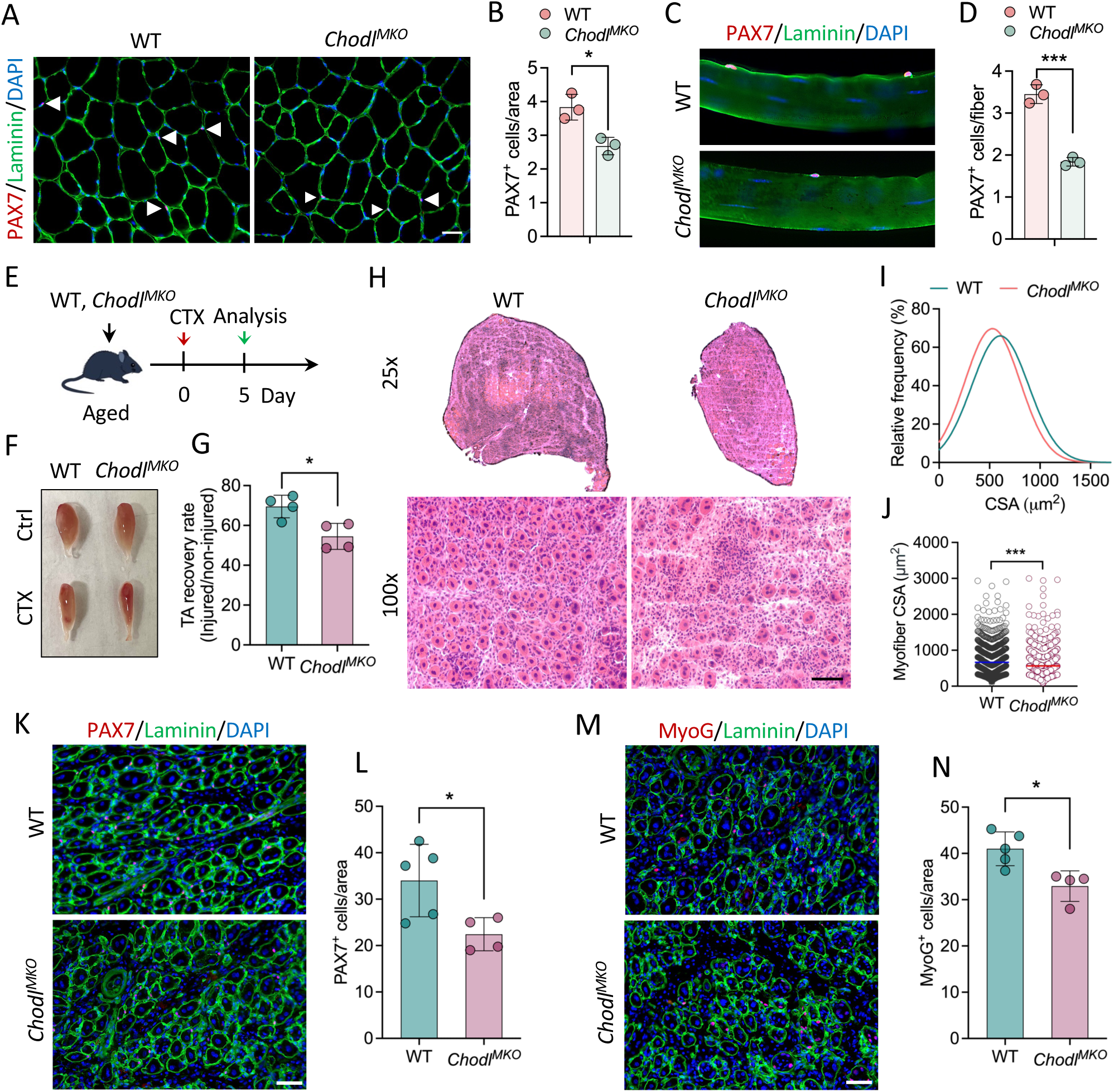
*Chodl* is necessary for SC maintenance and muscle regeneration in aging. **(A)** Immunofluorescence of PAX7, α-laminin on TA muscle cross-sections from old WT and *Chodl^MKO^* mice (22-month-old). Scale bar: 50 μm. **(B)** Quantification of PAX7^+^ cells per area as shown in (**A**), n=3 mice each group. **(C)** Immunofluorescence of PAX7, α-laminin on freshly isolated myofibers from old WT and *Chodl^MKO^* mice extensor digitorum longus (EDL) muscle. Scale bar: 20 μm. **(D)** Quantification of PAX7^+^ cells on myofibers as shown in (**C**). n=3 mice each group, 20-25 fibers per mice. **(E)** Schematics showing experimental design involving cardiotoxin (CTX)-induced muscle regeneration on old mice (20 to 22-month-old). **(F)** Representative images of TA muscles at 5 dpi in old WT and *Chodl^MKO^* mice. **(G)** TA muscle recovery rate in old WT and *Chodl^MKO^* mice. WT, n=5 mice; *Chodl^MKO^*, n=4 mice. **(H)** H&E staining of TA muscle cross-sections of old WT and *Chodl^MKO^* mice at 5 dpi. Scale bar: 50 μm. **(I, J)** Relative frequency of myofiber CSA (**I**) and dot plot of individual myofiber CSA (**J**). n=4 mice each group. **(K, L)** Immunofluorescence of PAX7 and α-laminin (**K**) and quantification of PAX7^+^ cell number (**L**) on TA muscle cross-sections. WT, n=5 mice; *Chodl^MKO^*, n=4 mice. Scale bar: 50 μm. **(M, N)** Immunofluorescence of MyoG and α-laminin (**M**) and quantification of MyoG^+^ cell number (**N**) on TA muscle cross-sections. WT, n=5 mice; *Chodl^MKO^*, n=4 mice. Scale bar: 50 μm. Data are represented as mean ± SEM; Unpaired Student’s *t*-test; \**P*< 0.05, \*\**P* < 0.01, \*\*\**P* < 0.001.

To assess the role of CHODL in SC function and muscle regeneration in aged mice, we analyzed TA muscles at 5 dpi following CTX-induced muscle injury in the aged WT and *Chodl^MKO^* mice (**Fig. 4E**). Compared with aged WT mice, aged *Chodl^MKO^*mice exhibited a decrease in TA muscle size after injury with markedly reduced muscle recovery rate (**Fig. 4F, G**). Histological analyses revealed fewer centrally nucleated myofibers and greater infiltration of interstitial mononuclear cells in *Chodl^MKO^* TA muscles, indicating impaired muscle regeneration (**Fig. 4H**). Specifically, the distribution of myofiber CSA displayed a clear left-shift in *Chodl^MKO^* TA muscles, with a significant decrease of average CSA (**Fig. 4I, J**). Moreover, immunofluorescence for PAX7 revealed a 37% reduction in the number of PAX7^+^ cells per TA muscle cross-sectional area in aged *Chodl^MKO^* mice compared with aged WT mice (**Fig. 4K, L**). Additionally, we performed MyoG immunofluorescence of TA muscle cross sections to examine the number of differentiating SCs. A significant reduction in MyoG^+^ cells was observed in aged *Chodl^MKO^* mice, with a 20% decrease compared with aged WT mice (**Fig. 4M, N**). Together, these results suggest that deletion of *Chodl* in SCs reduces SC number, diminishes SC differentiation, and impairs muscle regenerative capacity in aged mice.

### Cell autonomous role of CHODL in SC proliferation during muscle regeneration

As the regenerative and SCs defects in the *Chodl^MKO^* mice may be due to concomitant loss of CHODL in both SCs and myofibers, we sought to specifically KO *Chodl* in SCs to examine the role of CHODL in these cells. To this end, we crossed *Pax7^CreER^* mice with *Chodl^flox/flox^*mice to generatd a tamoxifen (TMX)-inducible SC-specific *Chodl* KO (*Chodl^PKO^*) model. We then deleted *Chodl* in SCs of adult mice by TMX injection, followed by CTX-induced injury of TA muscles to assess muscle repair at 3.5 dpi (**Fig. 5A**). To evaluate SC proliferation *in vivo*, 5-Ethynyl-2’-deoxyuridine (EdU) was administrated to mice 8 hours before euthanasia via intraperitoneal (IP) injection to mark proliferating cells (**Fig. 5A**). Immunofluorescent staining of embryonic myosin heavy chain (eMyHC) revealed fewer eMyHC^+^ myofibers in *Chodl^PKO^* muscles compared to WT mice following muscle injury at 3.5 dpi, which was consistent to the observations in *Chodl^MKO^*mice suggesting the defect of regeneration (**Fig. 5B, C**). In addition, a significant reduction in the number of PAX7^+^ SCs was observed in *Chodl^PKO^* muscles compared to WT muscles (**Fig. 5D, E**). Furthermore, the percentage of EdU^+^ cells among the PAX7^+^ cells was significant lower in *Chodl^PKO^* muscles compared to WT muscles with 7.1 versus 13.1 (**Fig. 5F, G**), indicative of a reduction in proliferation. These results suggest an essential role of CHODL in maintaining regenerative capacity of adult SCs by promoting their proliferative activity.

**Figure 5.**
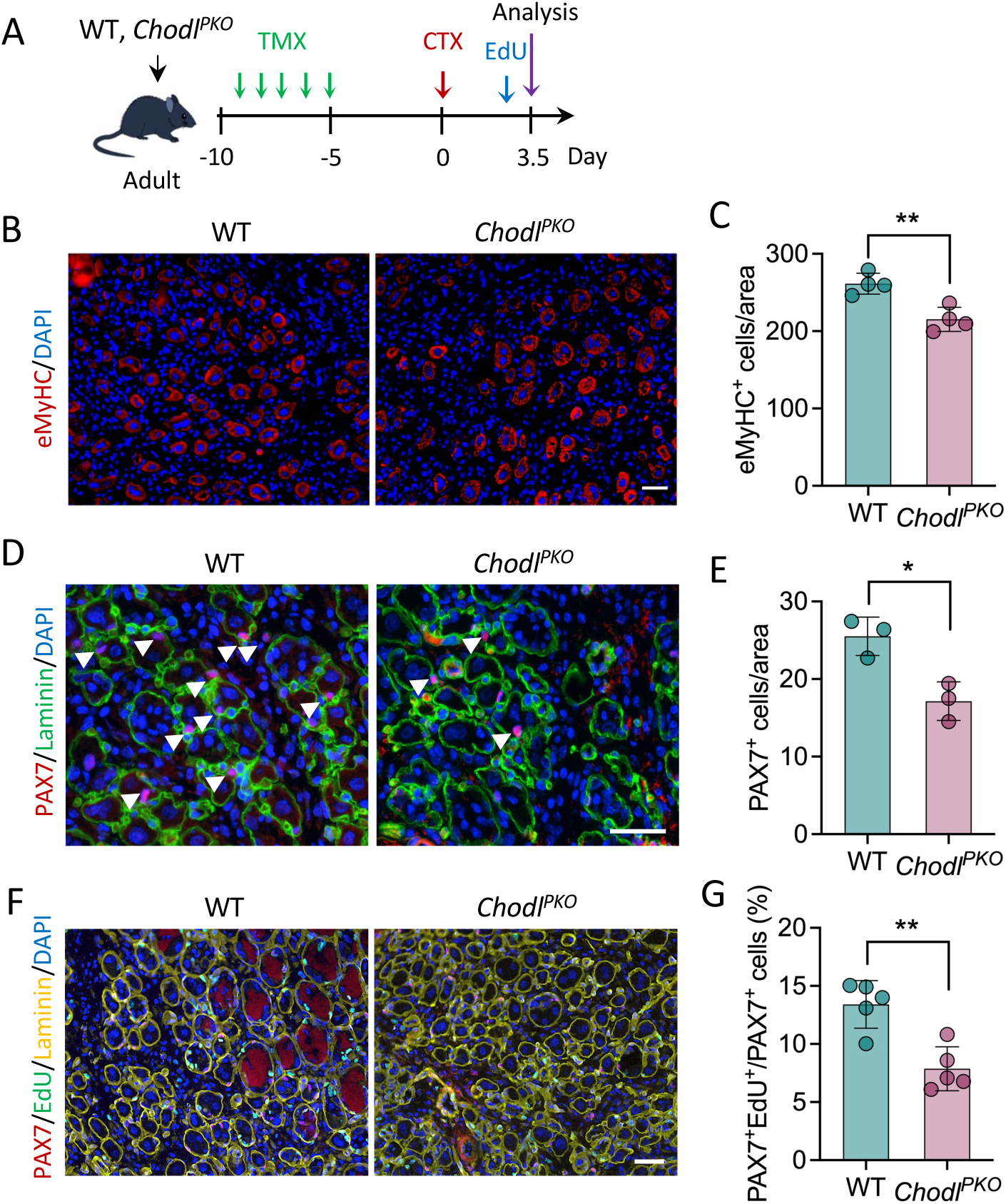
CHODL plays a cell autonomous role in adult SCs. **(A)** Schematics showing CTX injury and EdU labeling of cell proliferation in WT and *Chodl^PKO^* mice after tamoxifen (TMX) induction to knock out *Chodl* specifically in SCs. **(B)** Immunofluorescence of eMyHC on TA muscle cross-sections at 3.5 dpi. Scale bar: 50 μm. **(C)** Quantification of the eMyHC^+^ cell numbers per area as shown in (B). n=4 mice each group. **(D)** Immunofluorescence of PAX7 and α-laminin on TA muscle cross-sections at 3.5 dpi. Scale bar: 50 μm. **(E)** The average number of PAX7^+^ cells per area as shown in (**D**), n=3 mice each group. **(F)** Immunofluorescence of PAX7, α-laminin and EdU on TA muscle cross-sections at 3.5 dpi. Scale bar: 50 μm. **(G)** The average number of PAX7^+^/EdU^+^ and PAX7^+^ cells per area as shown in (**F**), n=5 mice each group. Data are represented as mean ± SEM; Unpaired Student’s *t*-test; \**P*< 0.05, \*\**P*< 0.01.

### *Chodl* KO affects expression of Notch signaling pathway and ECM component genes

To understand the molecular mechanism underlying CHODL function in SCs, we take advantages of scTenifoldKnk, a computational algorithm for predicting the molecular targets of genes of interests (Erbe et al., 2022). The scTenifoldKnk-based virtual KO analysis recapitulates most of the findings of real-animal KO experiments (Erbe et al., 2022; Osorio et al., 2022), validating its utility. To understand the molecular targets of CHODL in SCs, we obtained RNA-seq data from WT mice generated by Yue et al (Yue et al., 2017), and used the expression matrix from WT mice as the input for scTenifoldKnk. We constructed the WT GRN (Gene Regulatory Network) and then virtually knocked out *Chodl*. scTenifoldKnk analysis revealed 166 genes are differentially expressed between WT and *Chodl*-virtual KO with False Discovery rate (FDR) < 0.05 (**Fig. 7A**). More specifically, these virtual KO perturbed genes include a large cohort of extracellular matrix (ECM)-receptor interaction genes (*Col3a1*, *Col4a1*, *Col4a2*, *Col5a1*, *Col5a2*, *Col5a3*, *Col6a1*, *Cav1*, *Creb3l2*, *Erbb3*, *Fgfr1*, *Jsrp1*, *Lama2*, *Lamc1*, *Neb*, *Osm*), several myogenic marker genes (*Myf5*, *Myog*, *Mymk*) and Notch signaling pathway genes (*Notch3*, *Dll1*) (**Fig. 6A**). The perturbation of ECM genes suggests a key role of CHODL in SC niche function. Consistently, Notch signaling is not only crucial for maintenance of QSCs, but also plays a role in regulating SC niche (Brohl et al., 2012). The perturbation of myomaker (*Mymk*) in *Chodl* virtual KO is also consistent with its function in myoblast fusion (Millay et al., 2013).

**Figure 6.**
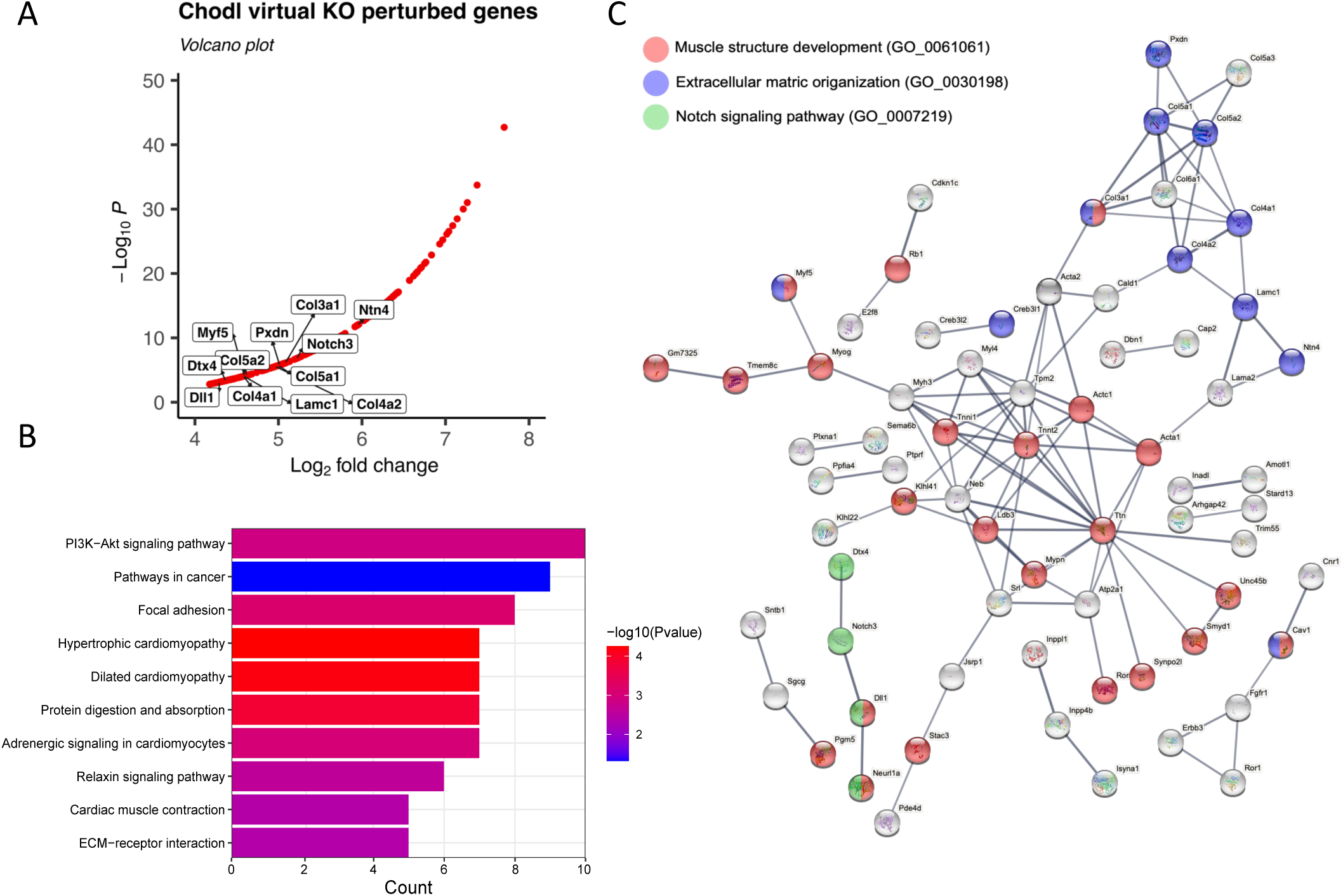
Virtual knockout analyses identify that ECM-receptor interaction is disturbed in *Chodl*-KO mice. scTenifoldKnk was used to achieve virtual knockout of *Chodl* in mouse skeletal muscle with the published RNA-seq data. By comparing the WT and pseudo-*Chodl* KO GRNs (Gene Regulatory Network), 166 genes were identified alterations in transcriptional regulatory networks and evaluates the knockout’s effects on the WT GRN. The resulting data was then used for enrichment analysis. **(A)** Volcano plot of *Chodl* virtual KO perturbed genes. **(B)** The KEGG pathway analysis of genes that are differentially expressed between WT and *Chodl-*virtual KO. **(C)** Interaction enrichment analysis of *Chod-*virtual KO perturbed genes.

**Figure 7.**
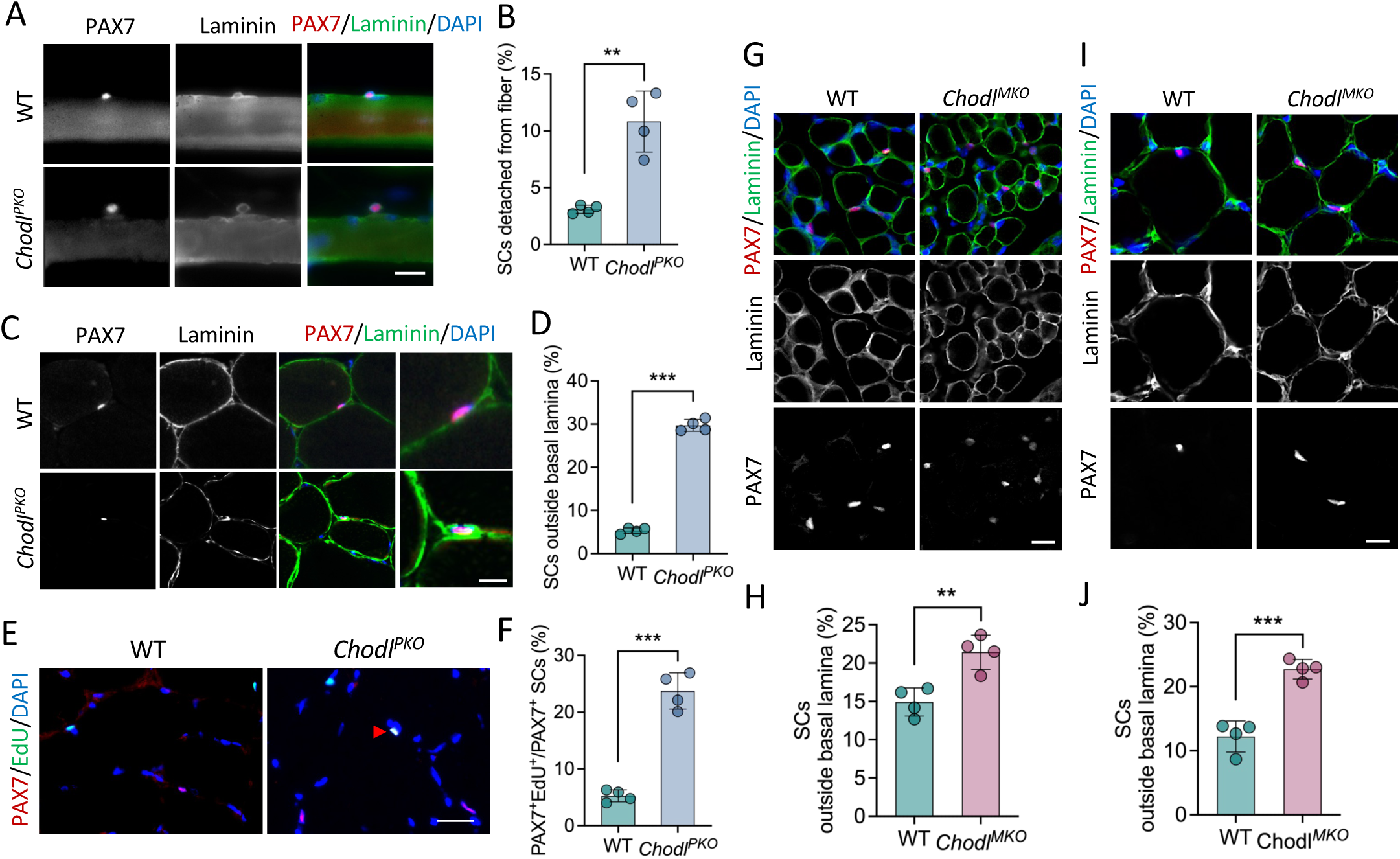
*Chodl* is critical for retaining SCs in myofiber niche during development. **(A)** Immunofluorescence of PAX7, α-laminin on freshly isolated myofibers from young adult WT and *Chodl^PKO^* mice extensor digitorum longus (EDL) muscle. Scale bar: 20 μm. **(B)** Percentage of SCs detached from fiber as shown in (**A**). n=4 mice each group, 20-25 fibers per mice. **(C)** Immunofluorescence of PAX7, α-laminin on TA muscle cross-sections of adult WT and *Chodl^PKO^* mice. Scale bar: 10 μm. **(D)** Percentage of SCs localized at interstitial space as shown in (**C**), n=4 mice each group. **(E)** Immunofluorescence of PAX7, EdU on TA muscle cross-sections of adult WT and *Chodl^PKO^* mice. Red arrowhead indicates EdU/PAX7 double positive cells. Scale bar: 20 μm. **(F)** Percentage of EdU^+^ to total PAX7^+^ cells as shown in (**E**), n=4 mice each group. **(G)** Immunofluorescence of PAX7, α-laminin on TA muscle cross-sections of WT and *Chodl^MKO^* mice at postnatal day 7 (P7). Scale bar: 50 μm. **(H)** Percentage of SCs localized at interstitial space as shown in (**G**), n=4 mice each group. **(I)** Immunofluorescence of PAX7, α-laminin on TA muscle cross-sections of WT and *Chodl^MKO^* mice at P21. Scale bar: 50 μm. **(J)** Percentage of SCs localized at interstitial space as shown in (**J**), n=4 mice each group. Data are represented as mean ± SEM; Unpaired Student’s *t*-test; ** *P*< 0.01, *** *P*< 0.001.

We further performed KEGG (Kyoto Encyclopedia of Genes and Genomes) pathway analysis of the 166 differentially expressed genes (DEGs). The results reveals that the DEGs were highly enriched in several pathways, including ‘PI3K-Akt signaling pathway’, ‘Focal adhesion’, ‘Cardiac muscle contraction’ and ‘extracellular matrix-receptor interaction’ (**Fig. 6B**). The results are consistent with a potential role of CHODL in mediating ECM interaction. Next, we applied the interaction enrichment analysis (Szklarczyk et al., 2015), which was based on the STRING protein-protein interaction database, on these 166 DEGs. We found that most of the DEGs appeared in a fully connected component in the STRING interaction network (**Fig. 6C**), indicating a tightly associated relationship between these genes. The core of the interactome map contains mostly muscle structural and metabolic genes (marked by red dots), while the ECM (blue dots) and Notch signaling (green dots) genes are clustered around the core genes (**Fig. 6C**). The in-silico analysis together indicates potential roles of CHODL in muscle structure (myofiber) and ECM interaction.

### CHODL is critical for retaining MuSCs in myofiber niche

Inspired by the potential role of CHODL in mediating ECM interaction that is crucial for SC maintenance affected by the *Chodl* KO, we investigated if CHODL plays a role in localization of SCs in the niche. We first isolated single myofiber from EDL muscles and label SCs with PAX7 and ECM with laminin (**Fig. 7A**). We observed that a substantial fraction of PAX7^+^ SCs (∼11%) was loosely associated with myofibers in *Chodl^PKO^* while only ∼3% of SCs were not closely attached to WT myofibers (**Fig. 7A, B**). To exclude the potential effects of collagen digestion during myofiber isolation, we examined SCs in cross sections of TA muscles of adult mice (**Fig. 7C**). In the WT muscle, SCs are predominantly located under the basal lamina (ECM), with only ∼5% located outside the basal lamina (**Fig. 7D**). In contrast, ∼30% SCs were found outside the basal lamina in the interstitial space in the *Chodl* KO muscles (**Fig. 7D**). Stem cell niche is crucial for maintaining stem cell quiescence. We further investigated whether the SCs in the interstitial space were precociously activated based on EdU uptake (**Fig. 7E**). We found that the percentage of EdU^+^ SCs in *Chodl^PKO^* mice was 5 times of that in WT mice (**Fig. 7F**). These results indicate that loss of CHODL disrupts sublaminar localization of SCs, inducing their precocious activation.

The interstitial localization of SCs in adult muscles could be due to their inability to maintain niche localization or defects in niche homing during development. A proportion fetal PAX7^+^ myoblasts took sublaminar position to become QSCs starting from fetal development throughout early postnatal growth (Kassar-Duchossoy et al., 2005; Schultz, 1978; Tamaki et al., 2002). This provides a time window to study homing of SCs. To evaluate the impact of *Chodl* KO on homing of SCs during muscle growth, we stained PAX7 and α-laminin to examine the localization of PAX7^+^ SCs at postnatal day 7 and 21 (P7 and P21) (**Fig. 7G-J**). Analysis of PAX7^+^ cells on TA muscles of WT and *Chodl^MKO^* mice reveals two phenomena. First, higher percentages of PAX7^+^ SCs were located outside the basal lamina in the *Chodl^MKO^* muscles than in the WT muscles, at both P7 and P21 compared to WT SCs (**Fig. 7H, J**). The difference in sublaminar SCs at such early stages is suggestive of homing defects instead of inability to maintain SC niche. Second, the percentages of interstitial SCs went down from 15% at P7 to 12% at P21 in the WT muscles (**Fig. 7H, J**), indicative of ongoing homing of SCs during postnatal growth and development. However, the percentages of interstitial extra-niche SCs remained at around 22% at both P7 and P21 (**Fig. 7H, J**), suggesting that homing is stalled during this period. These results together demonstrate that *Chodl* deletion profoundly affect homing of SCs during muscle development.

## Discussion

CHODL was initially identified as a member of the C-type lectins, and subsequent studies have shown that it is predominantly expressed in the vascular muscles of testes, muscle cells of prostate stroma, heart, and skeletal muscle (Claessens et al., 2007; Weng et al., 2003). The observation of highly enriched *Chodl* expression in quiescent SCs led us to hypothesize that CHODL might play a crucial role in regulating muscle stem cell function. In this study, we utilized two cell type-specific *Chodl* KO mouse models to investigate its function and demonstrated its essential role in SC pool maintenance and behavior, and muscle regeneration. Our findings indicate that loss of CHODL disrupts SC differentiation and impairs to maintain SC pool, which resulted in impaired muscle regeneration capacity post injury. Taken together, our study revealed a previously unrecognized role of CHODL in SC biology and its contribution to muscle regenerative capacity.

Despite the demonstrated importance of CHODL in SCs, its cellular mechanisms of action of CHODL remain poorly understood. CHODL has been shown to interact with ECM via its extracellular C-type lectin domain and with the Rab GTPase complex intracellularly, suggesting that it may mediate ECM signaling in SCs (Claessens et al., 2008; Oprisoreanu et al., 2019). We observed a significant delay in muscle regeneration following SC-specific *Chodl* deletion, accompanied by an increased number of *Chodl* KO SCs localized in the interstitial space of myofibers, which is indicative of SC activation. Ex vivo myofiber isolation further confirmed these findings, revealing a higher number of detached SCs in the absence of CHODL. This observation aligns with the proposed role of CHODL in ECM signaling, which is known to be critical for SC attachment to the myofiber surface (Gattazzo et al., 2014; Thomas et al., 2015; Urciuolo et al., 2013). Thus, the loss of CHODL may impair ECM signaling, leading to SC detachment and subsequent alterations in SC fate.

Maintaining the balance between quiescence and activation is a key function of the progenitor cell niche, with ECM interactions playing a pivotal role in regulating SC fate across various stem cell niches (Brunet et al., 2023; Fuchs and Blau, 2020; Hicks and Pyle, 2023; Thomas et al., 2015). Besides its structural function, the ECM acts as a signaling platform, where cell surface receptors interact with ECM proteins to regulate cellular behavior (Thomas et al., 2015). Notably, SCs cultured on ECM-coated plates (e.g., laminin and collagen) exhibit increased Pax7 expression compared to those grown on uncoated plates (Grefte et al., 2012; Wilschut et al., 2010). The Notch signaling pathway is a critical regulator of SC quiescence (Bjornson et al., 2012; Conboy and Rando, 2002), acting as a sensor of homeostatic conditions and reinforcing the niche by promoting the production of active collagen V, which in turn maintains SC quiescence (Baghdadi et al., 2018). A previous microarray study suggested that CHODL may contribute to basal lamina assembly through Notch signaling (Brohl et al., 2012). However, further investigation is still needed to delineate the precise signaling mechanisms by which CHODL regulates SC function.

Given the essential role of CHODL in SC maintenance and muscle homeostasis, it is important to explore its pathophysiological roles in various conditions. In experimental CTX-induced muscle injury, the initial decrease in CHODL expression may result from SC differentiation into myofibers during regeneration. However, it appeared that CHODL levels appeared to be restored post-injury to maintain an adequate SC pool. Our data suggest that this process depends on functional CHODL, as *Chodl* KO significantly reduced SC pool during injury recovery. Moreover, *Chodl* KO also led to a reduced the SC pool in aged mice. Previous studies have demonstrated a progressive decline in the number of SCs with aging, leading to impaired muscle regeneration and contributing to age-related functional decline (Blau et al., 2015; Chakkalakal et al., 2012). This depletion of SCs is a causative factor in the onset of various muscle disorders such as sarcopenia (Shefer et al., 2006). In aging muscle, the loss of SCs results in diminished regenerative capacity, primarily due to dysregulation of SC homeostasis and a reduction in self-renewal capacity (Brack and Munoz-Canoves, 2016; Brunet et al., 2023; Feige et al., 2018; Rando et al., 2025). Therefore, defining the role of CHODL in aging-related muscle decline and elucidating the signaling pathways it mediates may uncover novel therapeutic targets for preserving muscle function during aging.

In summary, our study highlights a critical role for CHODL in maintaining the SC pool, thereby supporting muscle homeostasis and post-injury regeneration. Future research aimed at delineating CHODL-mediating cellular pathways in regulating SC functions may provide insights into its pathogenic role and therapeutic potential in pathological conditions associated with muscle functional decline.

## Methods

### scRNA-Seq data analysis

scRNA-seq datasets of muscle satellite cell during muscle regeneration were downloaded from publicly available data (GSE138826) (Oprescu et al., 2020b). For data analysis, barcodes and reads were aligned to mm10 (*Mus Musculus*) using CellRanger v3.1, and data analysis was performed using Seurat v3.1 as previously described (Oprescu et al., 2020a).

### Mice

All procedures involving animals were conducted in compliance with National Institutes of Health and Institutional guidelines with approval by the Purdue Animal Care and Use Committee. *Chodl^fl/+^* mice were created via CRISPR/Cas9 technology. Firstly, two sgRNAs-targeting the introns on both sides of the floxed region (contains exons 2-4) of *Chodl* were synthesized and transcribed, respectively. The donor vector with the loxP fragment was designed and constructed in vitro. Then Cas9 mRNA, sgRNA and donor were co-injected into zygotes. Thereafter, the zygotes were transferred into the oviduct of pseudo pregnant ICR females at 0.5 days post coitum, and F0 mice was born 19–21 days after transplantation. Finally, F0 mice were crossed with C57BL/6J mice to create heterozygous mice, which were used to produce homozygous *Chodl^fl/fl^*mice. *Pax7^CreER/+^* (stock #017763) and *MyoD^Cre/+^* (stock #014140) mice were purchased from the Jackson Lab. Male or female mice were used and always gender matched for each specific experiment. Mice were housed and maintained in the animal facility, with free access to standard rodent chow and water.

### *In vivo* treatment

Tamoxifen (cat# 57900, Calbiochem, USA) was dissolved in corn oil at a concentration of 10 mg/ml. Both *Pax7^CreER^*; *Chodl^f/f^* and *Chodl^f/f^* mice were administered tamoxifen at a concentration of 100 mg/kg per day for five continuous days by intraperitoneal injection. In continuous labelling experiments, 2’-Deoxy-5-ethynyluridine (EdU, cat# NE08701, Carbosynth, USA) was administrated uninterruptedly to mice through drinking water (0.3 mg/ml) at 8 h before sampling. Drinking bottles were protected from light.

### Muscle injury and regeneration

Muscle regeneration was induced by cardiotoxin (CTX, cat#217503, MilliporeSigma, USA) injection. Adult mice were first anesthetized using a ketamine-xylazine cocktail, and CTX was injected (50 μL of 10 μM solution) into the TA muscle. Muscles were then harvested at the indicated days post injection to assess the completion of regeneration and repair.

### Primary myoblast isolation, culture, and differentiation

Satellite-cell-derived primary myoblasts were isolated from hindlimb skeletal muscles of *Pax7^CreER^*; *Chodl^f/f^*mice at the age of 6–8 weeks as previously described. Muscles were minced and digested in collagenase, type I (cat# LS004196, Worthington, USA) and Dispase B (cat# 4942078001, Roche, USA) mixture. The digestion was stopped with growth medium (Ham’s F-10 Nutrient Mix medium supplemented with 20% FBS, 4 ng/ml basic FGF, and 1% penicillin–streptomycin). Cells were then filtered from debris, centrifuged, and cultured in growth medium on collagen-coated cell culture plates at 37°C, 5% CO_2_. For in vitro genetic deletion, *Pax7^CreER^*; *Chodl^f/f^* primary myoblasts were induced by 2 days of 4-OH tamoxifen (0.4 μM; cat# 479002, Calbiochem), and the primary myoblasts treated with vehicle were set as the control. For differentiation, primary myoblasts were seeded on BD Matrigel-coated cell culture plates and induced to differentiate in a low serum medium (DMEM (cat# 11965092, Gibco, USA) supplemented with 2% horse serum (cat# 26050088, Gibco, USA) and 1% penicillin-streptomycin).

### Single myofiber isolation and culture

Single myofibers were isolated from Extensor Digitorum Longus (EDL) muscles of adult mice as previously described (Pasut et al., 2013). The EDL muscle was removed from the hindlimb of the mouse and digested with 0.2% Collagenase type I for 60 minutes in shaking water bath at 37 °C. Single muscle fibers were obtained by gently triturating the digested muscle using a glass pipet in DMEM under a dissection microscope. Released single myofibers were then transferred and cultured in a horse serum-coated Petri dish (60-mm) in DMEM supplemented with 20% FBS, 4 ng/ml basic FGF, and 1% penicillin–streptomycin at 37 °C for indicated days. Fresh isolated and cultured myofibers were fixed immediately in 4% paraformaldehyde (PFA) for further analysis.

### Histology and immunofluorescence staining

Whole muscle tissues from WT, *Chodl^MKO^* and *Chodl^PKO^* mice were dissected and frozen immediately in an O.C.T. compound (cat# 4583, Sakura, USA). Frozen muscles were cross-sectioned (10 μm) using a Leica CM1850 cryostat. The slides were subjected to histological H&E staining or immunofluorescence staining. For immunofluorescence staining, cross-sections, single myofibers, or cultured cells were fixed in 4% PFA for 10 min, quenched with 100 mM glycine for 10 min, and incubated in blocking buffer (5% goat serum, 2% BSA, 0.1% Triton X-100, and 0.1% sodium azide in PBS) for 1 hr. Samples were then incubated with primary antibodies and then secondary antibodies and DAPI. All images were captured using a Leica DM 6000B microscope, and images for WT and KO samples were captured using identical parameters. All the images shown are representative results of at least three biological replicates. Primary and secondary antibodies used in this study were follows: Fiber Type I (MYH7, cat# BA-D8, DSHB, USA, 1:100), Fiber Type IIA (MYH2, SC-71, DSHB, USA, 1:100), Fiber Type IIB (Myh4, cat# BF-F3, DSHB, USA, 1:100), PAX7 (DSHB, USA, 1:10), eMyHC (MYH3, DSHB, USA, 1:100), MyoG (DSHB, USA, 1:1000), α-laminin (cat# L9393, MilliporeSigma, USA, 1:1000), MF20 (MYH1E, DSHB, USA, 1:100), Goat anti-Mouse IgG1, Alexa Fluor 568 (cat# A-21124, Thermo Fisher, USA, 1:1000), Goat anti-Mouse IgG2b, Alexa Fluor 647 (cat# A-211242, Thermo Fisher, USA, 1:1000), Goat anti-Mouse IgM, Alexa Fluor 488 (cat# A-21042, Thermo Fisher, USA, 1:1000).

### Protein extraction and western blot analysis

Total protein was extracted from homogenized muscle tissue using RIPA buffer containing 25 mM Tris-HCl (pH 8.0), 150 mM NaCl, 1mM EDTA, 0.5% NP-40, 0.5% sodium deoxycholate, and 0.1% SDS, supplemented with proteinase inhibitor (PI) and phenylmethylsulphonyl fluoride (PMSF). Protein concentration was measured by BCA protein assay. Proteins were separated by electrophoresis, transferred to polyvinylidene fluoride (PVDF) membrane, blocked with 5% fat-free milk for 1 h at room temperature and incubated with primary antibodies overnight at 4°C. Immunodetection was detected using enhanced chemiluminescence western blotting substrate (Santa Cruz Biotechnology) on a FluorChem R system (Proteinsimple). Primary and secondary antibodies used in this study were as follows: CHODL (cat#PA5-69993, Invitrogen, USA, 1:1,000), TUBULIN (cat# ab6046, Abcam, USA,1:5,000), HRP AffiniPure goat anti-Mouse IgG (cat# 115-035-03, Jackson ImunoResearch, USA, 1:10,000), HRP AffiniPure goat anti-Rabbit IgG (cat# 115-035-03, Jackson ImunoResearch, USA, 1:10,000).

### Total RNA extraction and qRT-PCR

Total RNA was extracted from cells and tissues using TRIzol reagent (Thermo Fisher Scientific) according to the manufacturer’s instruction. A total of 2 μg of total RNA was reversed transcribed with random primers, M-MLV reverse transcriptase and DTT. Real-time qPCR was carried out in a Roche Light cycler 480 PCR system with SYBR green master mix and gene-specific primers. Relative changes in gene expression were analyzed using the 2^−ΔΔCT^ method and normalized to β-actin. Primers used in this study were as follows (Sequence 5’ –> 3’): *β-actin* forward primer: GTCCCTCACCCTCCCAAAAG, *β-actin* reverse primer: GCTGCCTCAACACCTCAACCC. *Chodl* forward primer: AGCGGAGATGGCCAAACATC, *Chodl* reverse primer: TTCAGCGGGCTCTGTTGGAT.

### Virtual knockout of *Chodl*

scTenifoldKnk was used to achieve virtual knockout of *Chodl* in mouse skeletal muscle with the published RNA-seq data (Osorio et al., 2022; Yue et al., 2017). In brief, scTenifoldKnk uses a gene-by-cell count matrix from the wild-type (WT) sample as input. It first constructs a WT GRN (Gene Regulatory Network) based on this matrix and then simulates Chodl knockout by removing the *Chodl* gene from the WT GRN, resulting in a pseudo-*Chodl* KO GRN. A network comparison method is then applied to identify differentially regulated (DR) genes by comparing the WT and pseudo-*Chodl* KO scGRNs. These DR genes, also referred to as virtual KO perturbed genes, then be used for gene function enrichment analysis.

### Statistical analysis

All analyses were conducted with unpaired Student’s *t*-test, with a two-tail distribution. All experimental data are represented as mean ± SEM. Comparisons with p values < 0.05 were considered significant.

## Data availability statement

Data sharing is not applicable to this article as no datasets were generated or analyzed during the current study. The data that support the findings of this study are available on request from the corresponding author.

## Authors’ contributions

F.Y. and S.K., conceptualized the study. L.G. and K.H.K. completed experiments. S.O. and X.C. performed transcriptome analysis. L.G., K.H.K., Y.L., and J.R. analyzed data and completed statistical analysis. L.G. and Y.L. wrote the original manuscript, F.Y. and S.K. revised the manuscript.

## Acknowledgments

This work was supported by grants from the US National Institutes of Health R01AR078695 and R01DK132819 to S.K., and UF Start-ups funds to F.Y. The authors thank Jun Wu for technical assistance.

